# Multi-view Co-training for microRNA Prediction

**DOI:** 10.1101/620740

**Authors:** Mohsen Sheikh Hassani, James R. Green

## Abstract

MicroRNA (miRNA) are short, non-coding RNAs involved in cell regulation at post-transcriptional and translational levels. Numerous computational predictors of miRNA been developed that generally classify miRNA based on either sequence- or expression-based features. While these methods are highly effective, they require large labelled training data sets, which are often not available for many species. Simultaneously, emerging high-throughput wet-lab experimental procedures are producing large unlabelled data sets of genomic sequence and RNA expression profiles. Existing methods use supervised machine learning and are therefore unable to leverage these unlabelled data. In this paper, we design and develop a multi-view co-training approach for the classification of miRNA to maximize the utility of unlabelled training data by taking advantage of multiple views of the problem. Starting with only 10 labelled training data, co-training is shown to significantly increase classification accuracy of both sequence- and expression-based classifiers, without requiring any new labelled training data. After 11 iterations of co-training, the expression-based view of miRNA classification experiences an average increase in AUPRC of 15.81% over six species, compared to 11.90% for self-training and 4.84% for passive learning. Similar results are observed for sequence-based classifiers with increases of 46.47%, 39.53% and 29.43%, for co-training, self-training, and passive learning, respectively. The final co-trained sequence and expression-based classifiers are integrated into a final confidence-based classifier which shows improved performance compared to each individual view. This study represents the first application of multi-view co-training to miRNA prediction and shows great promise, particularly for understudied species with few available training data.

**Availability:** Code is available at https://github.com/GreenCUBIC/miRNA_MVCT. All datasets are publicly available with accession numbers listed in the manuscript.

## I. INTRODUCTION

Deoxyribonucleic acid (DNA) is a double helix molecule that encodes all the genetic information for a species. For this genetic information to be made usable by the cells, DNA is transcribed into Ribonucleic acid (RNA). MicroRNA (miRNA) are transcribed regions of DNA that, after some biological processes, ultimately form short non-coding RNAs typically in the range of 18 to 25 nucleotides. MicroRNA are involved in cell regulation at the post-transcriptional and translational levels through the degradation and translation inhibition of messenger RNA (mRNA). MicroRNA act on mRNA through a silencing mechanism and target a wide range of mammalian mRNA, affecting biological activities such as cell cycle control [1], biological development [2, 3], differentiation [4] and stress response [5–7]. Evidence suggests that approximately 60-90% of mRNA have the potential to be targeted by miRNA [8]. Recent studies have also suggested that miRNA may act as tumor suppressors in different cancers such as liver [9], colon [10], prostate [11] and bladder cancers [12].

The identification of novel miRNA requires an inter-disciplinary approach, in which both computational and experimental methods are involved. Through experimental approaches, a number of miRNA have already been identified for many distinct species, ranging from more than 2000 known human miRNA, to less than ten known mallard miRNA [13]. These known miRNA have been used as a basis to create a number of computational methods to predict new miRNA. Such computational methods use supervised machine learning to identify and abstract the patterns present in previously discovered miRNA. These models seek to identify new putative miRNA sequences exhibiting these patterns and classify arbitrary sequence windows as being either miRNA or non-miRNA. There are generally two different approaches to developing computational predictors of miRNA. Methods that examine the RNA sequence directly are referred to as *de novo* predictors, while methods that look for evidence of miRNA biogenesis processing within next generation sequencing (NGS) expression data are referred to as NGS- or expression-based methods [14]. Sequence- and expression-based techniques detect pre-miRNA sequences forming miRNA-like hairpins and classify them based on the presence or absence of sequence- or expression-based features, respectively.

A significant challenge in the field of miRNA prediction is that, for many species, there is a scarcity of labelled examples of known miRNA. This makes the task of miRNA classification a very difficult matter, particularly for newly sequenced species where there are few known miRNA exemplars available. According to miRbase [13], of the thousands of existing species, experimentally validated miRNA sequences are available for only 271 species. Approximately 30% of these species have 15 or fewer known miRNA sequences, meaning that in general most species have very few training exemplars available. The majority of current *de novo* and NGS-based miRNA prediction techniques use supervised learning methods for the detection of novel miRNA, thereby requiring a large database of known miRNA. In addition, these methods do not always achieve high accuracy, resulting in many sequences being falsely predicted to be miRNA. These false predictions represent a substantial loss of resources, as they lead to unnecessary experimental validation costs. Previous work has demonstrated that relying on training exemplars from other species leads to reduced classification performance for the target species [15]. Therefore, if we are to create more effective miRNA predictors, either new miRNA sequences must be identified, or a method must be designed to create reliable miRNA predictors from smaller numbers of known miRNA exemplars.

While the scarcity of labelled known miRNA data presents a challenge, the emerging abundance of unlabelled sequence and expression data represents a hitherto untapped opportunity for miRNA prediction. This prompted us to propose a new semi-supervised miRNA prediction technique that extracts maximal information from both the limited labelled miRNA data and the unlabelled data, therefore requiring very few known miRNA. We aim to not only minimize the number of labelled training exemplars required, but also to improve the performance of current prediction methods using fewer samples. In this way, we can create a more efficient method for the identification of novel miRNA, particularly for species with few known miRNA, while maximizing the return on investment for costly wet-lab validation experiments. As mentioned, all previous methods use supervised machine learning methods, which only make use of labelled instances for classification. Semi-supervised machine learning methods also make use of both labelled and unlabelled data for classification; such methods are designed to work in situations where we have a small number of known exemplars and a large body of unlabelled data.

We have recently reported on the first application of semi-supervised machine learning for miRNA prediction [16]. In that study, active learning is applied to guide the wet-lab experiments to iteratively increase the set of known miRNA within a species. Significant increases in miRNA prediction accuracy were observed using active learning; however, this approach presupposes the availability of wet-lab experiments for validating specific putative miRNA to augment the training set. We here explore the application of another method of semi-supervised machine learning: multi-view co-training [17]. This approach differs from active learning in that no additional labelled training data or wet-lab experiments are required. Instead, the training set is grown using high-confidence predictions from the classifiers themselves. Over-training is avoided by leveraging multiple views of the problem and applying co-training rather than self-learning. As with active learning, multi-view co-training aims to learn patterns not only from the few labelled exemplars, but also from the unlabelled data. The present study represents the first application of multi-view co-training to the problem of miRNA prediction.

Multi-view co-training has been applied to a range of pattern classification problems over the past few years, especially those in which labelled data is either rare or comes at a very high expense. The application of co-training has mostly been focused on areas such as natural-language processing and signal processing, where data have multiple views, such as text data and web data [18]. Applications of multi-view co-training to bioinformatics have been limited to prediction of protein function [19], prediction of breast cancer survivability [20], detection of mis-localized proteins in human cancers [21], gene expression classification [18], cancer sample classification [22] and phenotype prediction [23]. We anticipate that multi-view co-training is likely widely applicable to the field of bioinformatics, since the lack of labelled data is a concern in most bioinformatics problem domains.

We propose using multi-view co-training for miRNA prediction, which makes use of multiple views of the problem to create distinct classifiers – one for each view. Each classifier is initially trained on a small set of labelled samples and is applied to an unlabelled data set. The most confidently predicted unlabelled instances from each of these views are added to the training set without experimental validation, and this process is repeated multiple times. In multi-view co-training, we take advantage of the fact that the problem of miRNA prediction can be approached from two distinct views: sequence based *de novo* prediction or expression-based NGS prediction. This creates two views for classification, an expression-based view and a sequence-based view. It should be noted that both of these views have been previously applied to miRNA classification independently [24] and as an integrated feature set [25, 26]; however, multi-view co-training has yet to be explored in the field of miRNA prediction. By applying multi-view co-training, we leverage each view using the other to create iteratively more powerful classifiers. The significant advantage of co-training is that there is no need for collecting any new labelled data (i.e. we do not require any new costly wet-lab experiments).

In the following sections, we demonstrate that leveraging multiple views of the miRNA prediction problem can be used, through multi-view co-training, to create more accurate predictors without actually labelling any new data. Performance is compared to self-learning, where a single view’s classifier is reinforced with its own high-confidence predictions. Ultimately, the two views are combined using a confidence-based system to achieve a unified predictor more accurate than either view.

## II. METHODS

### A. Data Set Selection

To develop and evaluate miRNA predictors, we require datasets of “positive” sequences, known to be miRNA, and “negative” sequences, that meet the fundamental criteria to be miRNA [27], but are known to have other functions (i.e. pseudo-miRNA). To demonstrate the broad applicability of the approaches developed here, we use datasets from six distinct species: mouse, fruit fly, cow, horse, chicken, and human. For each species, we require five data sources: genomic data (UCSC genome browser database [28]), short RNA NGS expression data (NCBI GEO database [29]), a set of known miRNAs (miRbase - release 22 [13]), a set of known coding regions (Ensembl sequence database [30]), and a set of non-miRNA known functional non-coding RNA (*Rfam* [31]). The data source for each species is summarized in Table 1.

**Table 1.**
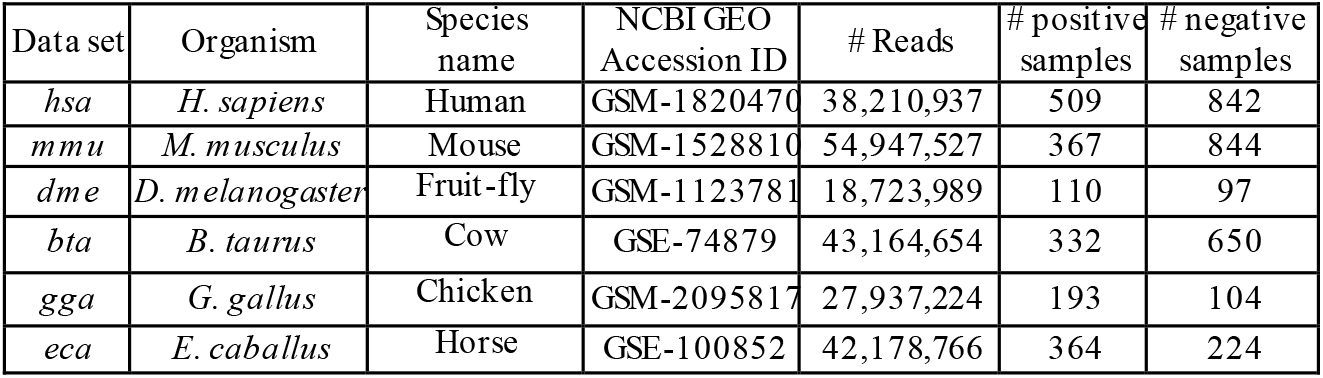
NGS data sets used in this study.

| Data set | Organism | Species name | NCBI GEO Accession ID | # Reads | # positive samples | # negative samples |
| --- | --- | --- | --- | --- | --- | --- |
| *hsa* | *H. sapiens* | Human | GSM-1820470 | 38,210,937 | 509 | 842 |
| *mmu* | *M. musculus* | Mouse | GSM-1528810 | 54,947,527 | 367 | 844 |
| *dme* | *D. melanogaster* | Fruit-fly | GSM-1123781 | 18,723,989 | 110 | 97 |
| *bta* | *B. taurus* | Cow | GSE-74879 | 43,164,654 | 332 | 650 |
| *gga* | *G. gallus* | Chicken | GSM-2095817 | 27,937,224 | 193 | 104 |
| *eca* | *E. caballus* | Horse | GSE-100852 | 42,178,766 | 364 | 224 |

Expression data were preprocessed using miRDeep2’s “mapper.pl” script [32] that automates the mapping of read stacks to the reference genome and the computation of RNA secondary structures for each mapped region. Given an NGS data set and a reference genome, this script produces a set of candidate pre-miRNA sequences accompanied by their structure and read stacks mapping to the sequence region. These candidate pre-miRNA sequences are compared to only the <u>high confidence</u> miRbase dataset for that species; those sequences that match to a known miRNA sequence are labelled as true positives. Remaining candidate pre-miRNA form a candidate negative set. These sequences are then aligned against the coding region data of the related species using *bowtie* [33] (up to two mismatches permitted). Since sequences known to form mRNA are unlikely to also form miRNA, all candidate negative sequences that align to a coding region are selected as our final true negative data. To ensure that the miRNA predictors developed here are not simply distinguishing between coding and non-coding regions, all available functional non-coding RNA within the 6 species were added to the negative set; these sequences were annotated in *Rfam* [31]. These supplementary negative sequences are functional non-coding RNA such as transfer RNAs (tRNA) and small nucleolar RNAs (snoRNA). Finally, *CD-HIT* [34] is applied (sequence identity < 90%) is used to remove redundant and highly similar sequences. Table 1 summarizes the final count of true positives and negatives for each of the species.

### B. Feature set selection

Data from all species were pooled for each of the sequence and expression-based views separately. Sequence-based features are obtained from the widely used HeteroMiRPred collection of available features [35]. These features include sequence-based, secondary-structure-based, base-pair-based, triplet-sequence-structure-based and structural-robustness-related features. To find the most informative sequence-based features, the correlation-based feature subset selection method in the Weka package [36] was applied using default parameters to all the training data from the six species. This algorithm minimizes the correlation between selected features while maximizing their predictive strength. This results in a vector of 32 sequenced-based features, pertaining to minimum free energy derived features, sequence/structure triplet features and dinucleotide sequence motifs, and structural robustness features. The eight expression-based features derived in [25] were used as our expression-based feature set, which consist of: (1) percentage of mature paired miRNA nts, (2) number of pairs in lower stem, (3) the percentage of RNA-seq reads in region which are inconsistent with Dicer processing and (4) from the loop region that match Dicer processing, (5) the percentage of RNA-seq reads - (6) RNA-seq-reads from the mature miRNA and - (7) RNA-seq-reads from miRNA* region which match Dicer processing, and (8) the total number of reads in the precursor region, normalized to experiment size.

### C. Classification Pipeline

All classifiers in this experiment are built using SKLearn random forest library [37], using default parameter values except for the number of trees, which was set to 500. Previous studies have demonstrated that random forest classifiers are effective for the prediction of miRNA [15, 38] and result in a balance between sensitivity and specificity [39–41]. We begin with a large set of labelled data. We then simulate the case where only a small subset of the data is labelled (forming our seed training set and independent test set), whereas the majority of data are simulated to be unlabelled. This represents a scenario for an understudied or newly sequenced species, where only a small number of known miRNA would be available with which to create the miRNA classifier.

For each species, the data are split into 20% hold-out data, which is used for testing the performance of the classifier, and the remaining 80% that is used for training the classifier. This distribution of data is randomly repeated 100 times, resulting in different seed training, test, and unlabelled sets. Therefore, each learning strategy described in this paper is evaluated 100 times using different initial seed training sets. The initial classifier for each view is built using a seed training set of 10 labelled samples (5 positive, 5 negative) randomly selected from the 80% split. After the initial model is built, it is applied to the remainder of the 80% “unlabelled” set, where the single most confident positive and negative predictions will be selected by each view. These two instances will be removed from the unlabelled set of their own view and added to the training set of the opposite view. The classifier for each view is re-trained using their new training sets, and this procedure is continued for several iterations. After each iteration, the learning curves of each view are plotted, where the area under the precision-recall curve (AUPR) is used as the summary performance metric over the 20% hold-out test dataset.

In the present study, for all six species, a total of 11 iterations were performed. More dynamic stopping criteria are available, including the approach suggested by Lewis and Gale [7], where the learning curve trends are monitored at each iteration; the learning process is stopped once maximum effectiveness has been reached and performing extra iterations does not result in a significant increase in performance. As mentioned above, in each iteration, the most confidently predicted positive and negative samples are added to the labelled training set of the alternate view. Clearly, more samples could be added per iteration to reduce computational load and expedite convergence. However, only two samples were selected in each iteration to ensure that only high-confidence predictions were being included in subsequent training sets. Our co-training pipeline can be observed in Figure 1.

**Figure 1.**
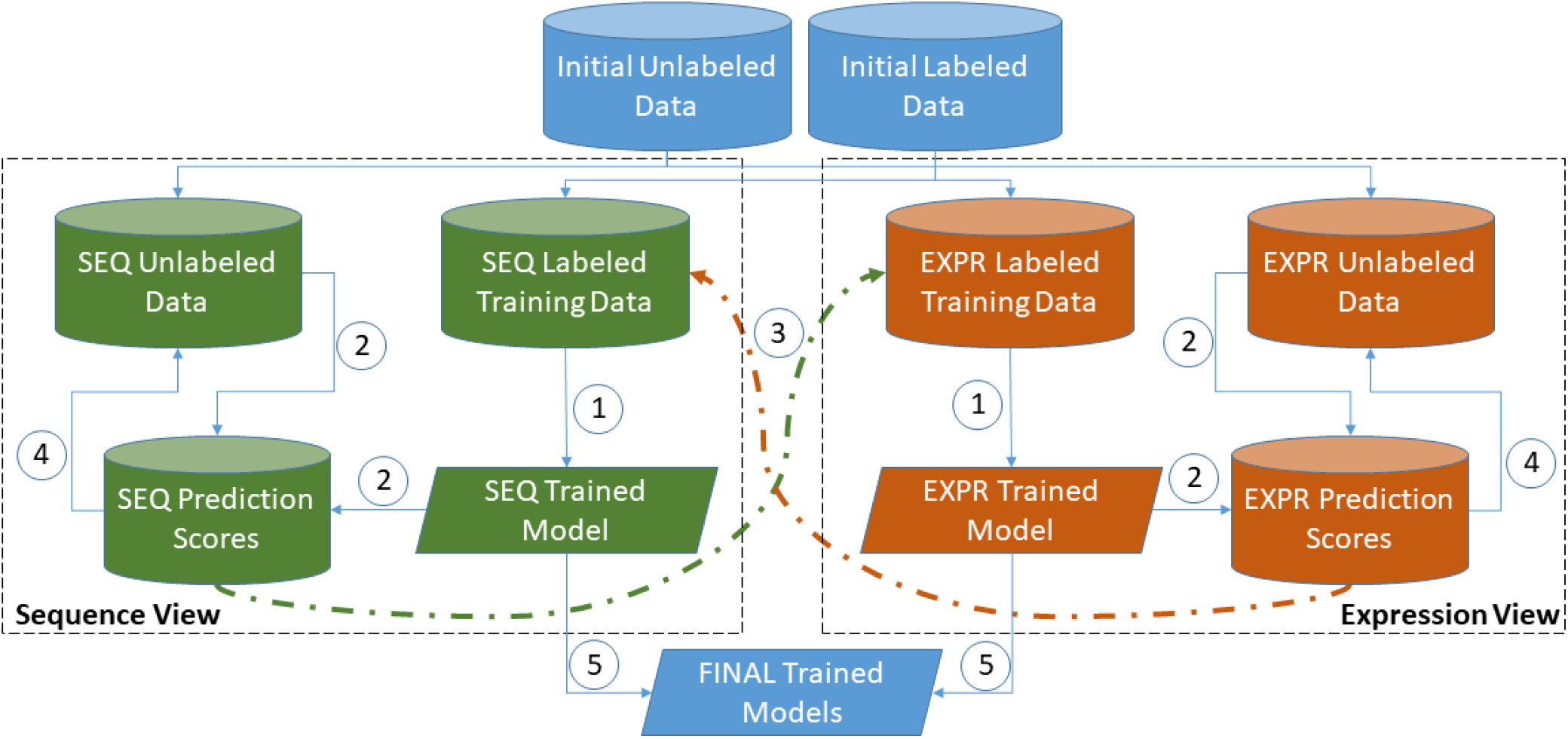
Multi-view co-training miRNA prediction framework. 1: Train Model. 2: Apply Model to unlabelled data to generate scores. 3: Add 2 top-scoring samples to other view’s training data. 4: Remove 2 top-scoring samples from unlabelled data. 5: Repeat steps 1-4 until stopping criterion; output final models.

As can be observed from Figure 1, we used a slightly different approach compared to the standard co-training approach originally proposed by Blum and Mitchell [17]. In most co-training models, a pool is created containing all the labelled data. At each iteration, the training set for each classifier is formed from a random subset of the total labelled pool. Also, at the end of each iteration, each classifier contributes newly labelled samples to that common pool. Our approach differs in that each classifier has its own independent labelled training data and, at the completion of each iteration, each classifier contributes newly labelled data (i.e. its high confidence predictions) to the other classifier’s training pool, not its own training set or to a shared pool.

### D. Comparing Co-training With Benchmark Algorithms

To evaluate the effectiveness of our proposed co-training method, we compared our results with two benchmark algorithms: a passive learning and self-training approach. The passive co-training strategy follows the multi-view co-training strategy outlined in Figure 1 exactly, except that the two samples selected to be added to the other view’s labelled training data were chosen randomly, rather than based on prediction confidence. In this approach, at each iteration the two random forest classifiers built using the two view-based feature sets and the training data were applied to the 80% unlabelled set. For each view, two randomly selected instances were removed from the unlabelled set and added to the training set of the other view using the predicted class labels. The selected instances were not selected completely at random; rather, they were selected randomly from a group of predictions expected to be either in the positive or negative class. To explain further, when the model was applied to the unlabelled set, each of the unlabelled instances received a prediction score. This score indicated the probability of that instance being in the positive or negative class, with a range of 0 to 100% probability for each class. In the passive approach, we randomly selected one instance that was more than 50% predicted to be a positive, and one that was more than 50% predicted to be a negative. In this way, the intention was to add one positive and one negative sample at each iteration to keep the training set of the passive approach in balance with that of co-training. As with the actual multi-view co-training strategy, this process was repeated for 11 iterations to enable direct comparison using the same number of training instances. By comparing our co-training classifier’s performance to that of this passive classifier, we can determine the significance of adding the most confidently predicted points in performance, while controlling for training set size.

In addition to passive co-training, self-training was used as another benchmark algorithm against which to compare our co-training approach. In self-training, each view was strengthened by its own predictions, meaning that learning was completed in each view independently [42–44]. In this approach, we created two classifiers, one for each of the sequence- and expression-based views. Each classifier was built using the same random forests, view-specific feature sets, and the same training, testing and unlabelled data sets previously described. Each classifier was built using its respective training set and, at each iteration, the trained model was applied to the unlabelled set. The single most confident positive and negative predictions were then removed from the unlabelled set and added to the training set of the view itself. This process was continued for the same number of iterations as co-training to fairly compare the results. By making this comparison, the value of using different views to strengthen each other can be observed. It was hypothesized that co-training would outperform self-training since using one’s own predictions to reinforce one’s beliefs is expected to lead to over-specialization (or drift) and reinforcement of a classifier’s errors. This is avoided in co-training since each classifier augments the training set of the other view, rather than its own.

### E. Combining the Views

So far, we have evaluated the performance of each of the individual views developed using multi-view co-training. Co-training is frequently used as a means to create a larger labelled data set, rather than to perform classification. However, if we were to use co-training itself as a classification tool, it would be better to combine both developed views at the end of their learning to produce a more robust classifier. Several ways can be thought of to combine the multiple views in co-training to arrive at a consensus decision. For the “classic” co-training approach [43], there is a single pool of quasi-labelled training data. Therefore, a final integrated classifier can be trained from that common pool, leveraging features from both views. In the cases where there are three or more views available for the problem, voting amongst the views for the class of the instance would be another reasonable approach. In our work, we create a combined co-training classifier considering the prediction confidence of both views for each instance. We do so using the prediction score produced by our algorithm for each instance and consider this a proxy for prediction confidence. When we obtain each views’ prediction score for an instance, we compare the two scores produced by each view, and give the preference to the view with the higher confidence in prediction. For example, if an instance is predicted by the sequence-based view to have a 70% probability of being a miRNA, while the expression-based classifier predicts it to have a 60% probability of not being a miRNA, the combined co-training classifier will predict the instance to be a miRNA.

## III. RESULTS

### A. Establishing the Effectiveness of Co-Training for miRNA Prediction

To demonstrate the effectiveness of applying multi-view co-training for miRNA prediction, we generated the performance learning curves of each view for the six species. Figure 2 illustrates the mean performance of the sequence and expression-based classifiers at each iteration of co-training for all data sets, averaged over 100 experiments with randomly selected seed training sets.

**Figure 2.**
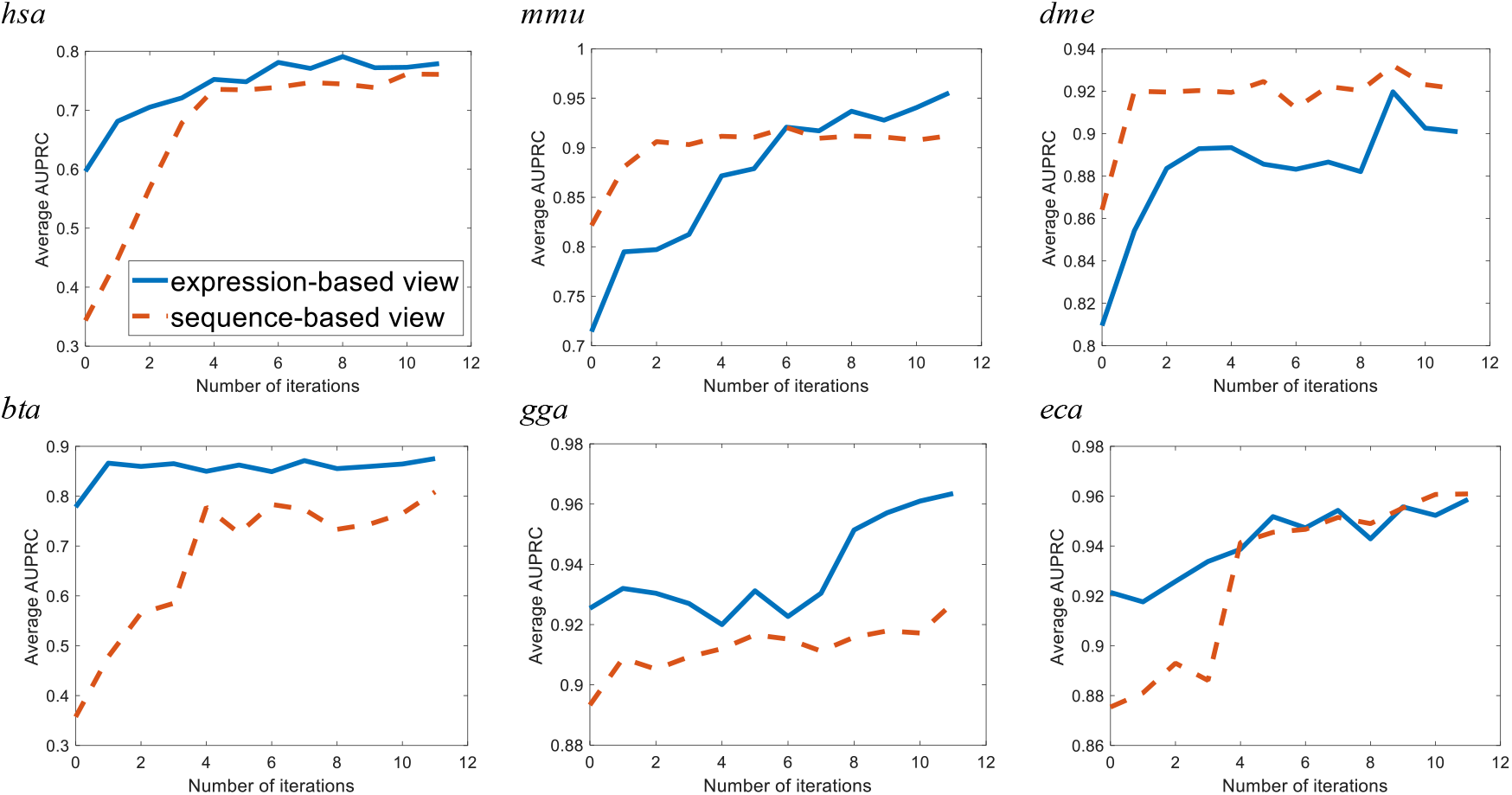
Performance of the sequence and expression-based classifiers in terms of average area under precision-recall curves at each iteration of co-training for all data sets over 11 iterations.

For all six species, substantial improvements in AUPRC are observed over the 11 iterations. The increase in performance is most dramatic in the human (*hsa*) and cow (*bta*) data sets. In human, the expression-based classifier’s average AUPRC increases by 30.6% and the sequence-based classifier more than doubles in average AUPRC, undergoing a 121% increase. Multi-view co-training appears to be least effective for chicken (*gga*) and horse (eca). The application of 11 iterations of co-training to the chicken data set resulted in the expression and sequence-based classifier’s average AUPRC increasing by 4.12% and 3.80%, respectively. When applied to the horse data set, the expression-based classifier’s average AUPRC is enhanced by 4.05%, where the sequence-based classifier undergoes an increase of 9.77%. However, for both chicken and horse, the initial classifier was already surprisingly effective with initial AUPRC values of over 0.87 for both, leaving little room for improvement due to co-training.

### B. Comparing Co-training With Benchmark Algorithms

Multi-view co-training was compared with two benchmark algorithms: 1) passive co-training, where samples are randomly selected at to be added to the training set of the alternate view in each iteration, and 2) single-view self-training, where learning is completed independently in each view and high confidence predictions are added to each view’s own training set at each iteration. In addition, the final iterations of co-training, passive learning and self-training are compared against a classifier without any iterative learning being applied (i.e. a classifier trained solely using the initial seed of 10 samples). This additional benchmark method is referred to as the “no-learning” classifier and corresponds to iteration zero in Figure 2. After 11 iterations, the final performance of all methods are plotted on a precision-recall curve for each species. The full set of PR-curves for all six species are included in Supplementary Figures 1 and 2. Table 2 summarizes these curves using mean area under precision-recall curve (AUPRC) over 100 random-seed repetitions. The final row of the table indicates the percent increase for each method relative to the no-learning classifier, averaged across all six species. For both the expression- and sequence-based classifiers, it can be observed that, in all six species, co-training outperforms all other methods. For clarity, the standard deviations are not shown, which were in the range of 0.001 to 0.003 for the different experiments. To test for the statistical significance of differences between methods observed in our results, the ANOVA test was first applied to the results in Table 2. This test indicated that there is a statistically significant difference between the mean results of the groups (α = 0.05). The Tukey test was then applied between the co-training results and each of the other methods, and all results were found to be statistically significant (at α =0.01).

**Table 2.** Final AUPRC classification results using co-training, single-view (self-training), passive learning, and no-learning for expression and sequence-based views. Values in parentheses represent the percent increase relative to the no-learning classifier. The last row indicates average increase compared to the base “no-learning” classifier, averaged over all six species.

| Data set | Expression-based classifier’s average AUPRC |  |  |  | Sequence-based classifier’s average AUPRC |  |  |  |
| --- | --- | --- | --- | --- | --- | --- | --- | --- |
|  | No-learning | Passive learning | Self-training | Co-training | No-learning | Passive learning | Self-training | Co-training |
| *hsa* | 0.597 | 0.672(+12.7%) | 0.770(+29.0%) | 0.779(+30.6%) | 0.344 | 0.616(+79.3%) | 0.720(+109.66%) | 0.761(+121.4%) |
| *mmu* | 0.714 | 0.708(-0.83%) | 0.883(+23.7%) | 0.955(+33.7%) | 0.822 | 0.887(+7.91%) | 0.895(+8.87%) | 0.912(+11.0%) |
| *dme* | 0.810 | 0.883(+9.10%) | 0.886(+9.50%) | 0.901(+11.3%) | 0.864 | 0.909(+5.16%) | 0.912(+5.51%) | 0.921(+6.6%) |
| *bta* | 0.778 | 0.827(+6.23%) | 0.815(+4.70%) | 0.865(+11.1%) | 0.357 | 0.654(+83.1%) | 0.732(+104.7%) | 0.809(+126.3%) |
| *gga* | 0.925 | 0.923(-0.22%) | 0.946(+2.26%) | 0.964(+4.12%) | 0.893 | 0.894(+0.08%) | 0.911(+2.03%) | 0.927(+3.79%) |
| *eca* | 0.921 | 0.941(+2.11%) | 0.942(+2.25%) | 0.958(+4.02%) | 0.875 | 0.884(+0.99%) | 0.932(+6.43%) | 0.961(+9.74%) |
| **Avg.** | - | **+4.84%** | **+11.90%** | **+15.81%** | - | **+29.43%** | **+39.53%** | **+46.47%** |

By comparing our multi-view co-training approach against single-view self-training and passive approaches, we can better comprehend the value of using two views and adding the most confidently predicted instances to the training set. The passive co-training approach serves as a control for training set size, whereas the self-training approach highlights the value in using two independent views of the problem. From the AUPRC values in Table 2, it is evident that co-training outperforms the other classifiers over all of the six data sets. By demonstrating substantially better results than the passive learning and self-training classifiers, it can be concluded that miRNA classification using co-training is the superior method.

### C. A Combined Co-training Classifier

We apply our proposed combined co-training algorithm to the data we had previously gathered for the individual views. The performance of the expression and sequence-based classifiers and the combined co-training approaches are compared in Figure 3 in terms of the average AUPRC values. These results illustrate the mean of 100 experiments with random seeds, as previously mentioned. In all cases, the combined co-training shows enhanced performance compared to each of the individual views on all data sets by taking advantage of the results from the dominant view.

**Figure 3.**
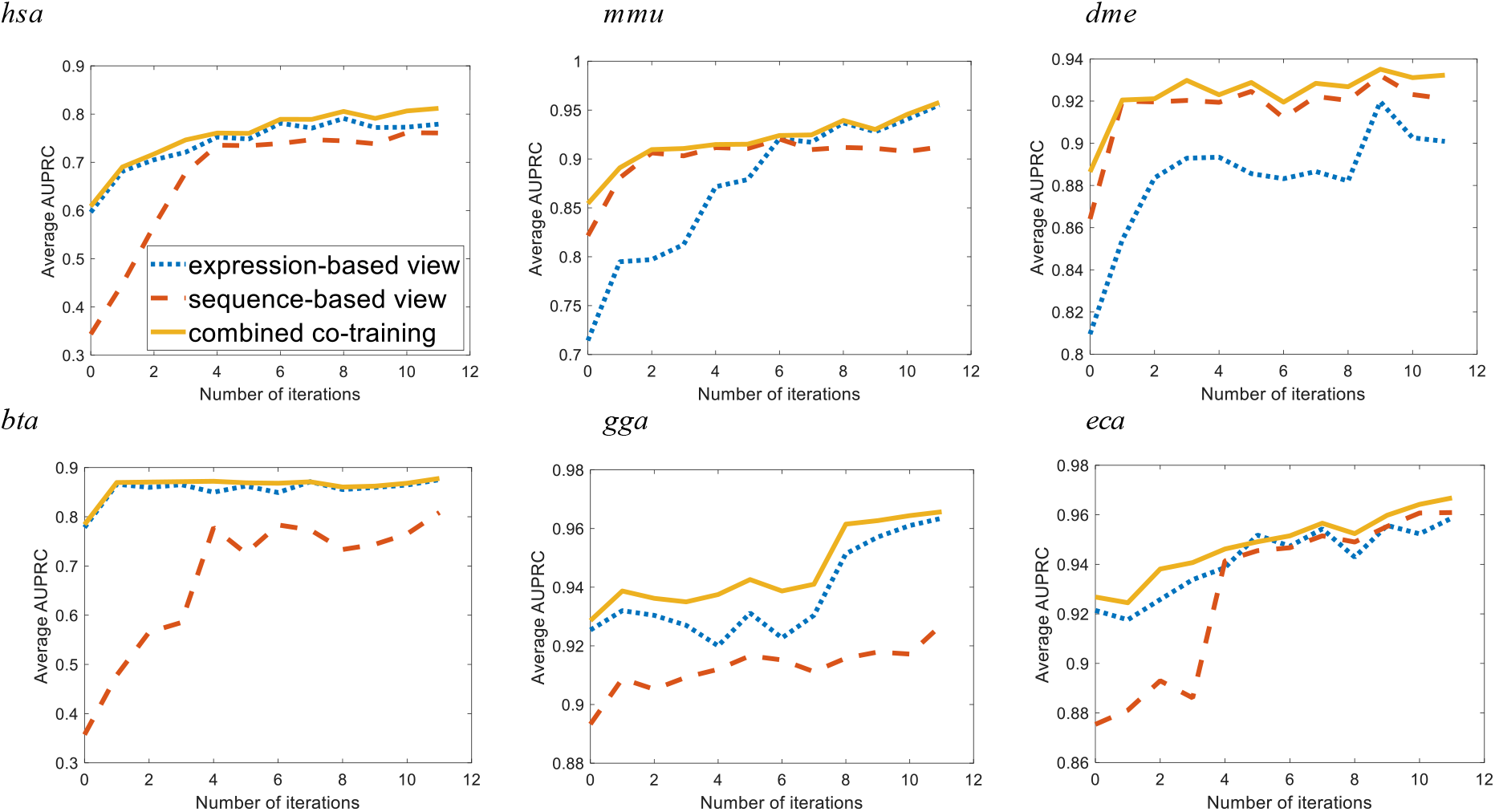
Performance of the sequence and expression-based classifiers and the combined co-training classifier in terms of average area under precision-recall curves at each iteration for all six data sets over 11 iterations.

## IV. DISCUSSION

During our experiments, we compared multi-view co-training to self-training. In the self-training approach, although the most confidently predicted points are added to the training set at each iteration, we only make use of one view of the problem. By making this comparison to our multi-view co-training approach, we are observing the effectiveness of strengthening one view using the other. By comparing the results of self-training to our co-training approach in Table 2, the advantage of using multiple views for training become clear. Here we observe an average co-training AUPRC that is 3.91% and 6.94% higher than that of the self-training classifier over all the data sets for expression- and sequence-based classifiers, respectively. These results highlight the significance of using multiple views (where applicable) for solving a pattern classification problem.

When examining the learning curves for self-training (data not shown), it was observed that in some cases the performance increase would plateau at a much earlier stage compared to co-training. This may be due to the fact that the amount of evidence that one view can provide is limited. In these cases, a single-view classifier quickly exhausts the information available from that view. Adding any high confidence instances predicted by such a classifier to the training set will not further strengthen the view; beyond this point, the classifier is simply adding instances with similar characteristics to those already in its labelled training set, therefore providing the classifier with redundant data. In contrast, using a different view for augmenting the training set could be highly valuable.

In the passive co-training approach implemented in this study, despite making use of both views of the problem similar to the co-training approach, we do not use the most confidently predicted instances. Instead, we choose expected positive and negative instances randomly and without knowing the model’s confidence in their labelling. The results of this approach show a high degree of fluctuation during iterations, demonstrating performance increase in some and decrease in the others. This is due to the random selection of instances to add to the training set, which in some cases leads to a mislabelled instance being added to the training set, therefore decreasing the classifier’s performance. When comparing the results of this passive approach to our co-training approach, it can be observed that strictly adding high confidence predictions has a significant effect on the classifier’s performance. By doing so, co-training demonstrates a 10.97% higher average AUPRC over all the data sets for the expression-based classifier, and a 17.24% higher average AUPRC for the sequence-based classifier. In addition, it can be observed that for the expression-based classifier in the chicken and mouse data sets, passive learning actually decreased the performance of the original models by 0.22% and 0.83%, respectively. This means that for these experiments, the addition of 22 instances resulted in weaker classifiers rather than strengthening them. This is due to the fact that a number of mislabelled points have been added to the original training sets for these species, causing the models to become weaker, since they are trained using incorrectly labelled data. These results underlie the importance of strictly adding high confidence instances to the training set, and that using multiple views alone is not sufficient for building a robust classifier.

The advantage of using multi-view co-training can be clearly seen in cases where one view initially performs much worse than the other. From Figure 2 it can be observed that in all of the data sets, one view clearly outperforms the other before the start of co-training. After 11 iterations, it is observed that the weaker view greatly improves in performance, and approaches the performance of the stronger view, even surpassing it in two data sets (mouse and horse). In other words, by using multiple views, one view can compensate for the weak performance of another view. It must also be mentioned that during the different iterations of co-training, no misclassified instances were added to the training set of neither view. This means that in all cases, the top prediction of both views was a correct prediction, demonstrating the accuracy of the classifiers developed using co-training. The fact that all instances that are newly added to the training set are correctly predicted is evidence that this approach is successful.

The co-training variation we have implemented in this study (explained in section C of Methods) was first proposed by Brefeld and Scheffer [45] and has been since used in [19]. This variation of multi-view co-training ensures that the views do not converge and that they are learning from different instances, since the majority of the training sets from each view are non-overlapping. Therefore, this method will further strengthen each view by directly incorporating evidence provided by another view into its training data. Since in this method each algorithm learns from the top predictions of the other algorithm, the algorithm is exposed to the risk of drift. That is, if one view starts making false predictions, these will quickly be amplified. To reduce the chances of this occurring we only allow each classifier to add one positive and one negative instance to the training set of the other classifier at each learning iteration. Future work could compare this approach to traditional co-training, where a single labelled training set is used, to compare their robustness to drift over a number of iterations. Since we ultimately require a single classification from both views, we combined the final view-based classifiers trained using co-training to create a combined co-training classifier. The proposed confidence-based combination rule proved to be highly effective. This indicates that mispredictions from individual views tend to have lower confidence than the correct predictions from the alternate view. By incorporating both views, we can take advantage of a greater number of strong predictions, therefore minimizing the number of weak predictions in our classifier and increasing performance.

## V. CONCLUSION

In this study, we propose a novel multi-view co-training approach for the classification of miRNA. By using a multi-view approach, we take advantage of both sequence and expression-based features to maximize miRNA classification accuracy, as measured using AUPRC. Using 11 iterations of co-training, the expression-based view of miRNA classification experiences a 15.81% increase on average over all data sets, compared to 11.90% for self-training and 4.84% for passive learning. The sequence-based classifier also experiences an increase of 46.47% across all species on average, where self-training and passive learning show 39.53% and 29.43% improvements, respectively. Finally, the developed sequence and expression - based classifiers are integrated into a final confidence-based combined co-training classifier which shows improved performance compared to each of the individual views. Considering the fact that we are using an absolute minimum number of labelled instances for classification (5 positive, 5 negative), these results demonstrate that multi-view co-training is a highly effective approach for miRNA classification. This approach is expected to be particularly useful in cases where labelled training is scarce, such as for newly sequenced species that have yet to be thoroughly analyzed or annotated through costly web-lab experiments.

## ADDITIONAL INFORMATION

### Ethics approval and consent to participate

Not applicable

### Consent for publication

Not applicable

### Availability of data and material

Our method is available at https://github.com/GreenCUBIC/miRNA_MVCT. All datasets are publicly available with accession numbers listed in the manuscript.

### Authors’ contributions

JG conceived of the study. MSH developed the algorithms and conducted the experiments. MHS wrote the initial draft and both authors reviewed and approved of the final manuscript.

